# Multiview Graph Learning for single-cell RNA sequencing data

**DOI:** 10.1101/2021.11.05.467476

**Authors:** Abdullah Karaaslanli, Satabdi Saha, Selin Aviyente, Tapabrata Maiti

**Author notes:** To whom correspondence should be addressed. Equal contribution.

## Abstract

Characterizing the underlying topology of gene regulatory networks is one of the fundamental problems of systems biology. Ongoing developments in high throughput sequencing technologies has made it possible to capture the expression of thousands of genes at the single cell resolution. However, inherent cellular heterogeneity and high sparsity of the single cell datasets render void the application of regular Gaussian assumptions for constructing gene regulatory networks. Additionally, most algorithms aimed at single cell gene regulatory network reconstruction, estimate a single network ignoring group-level (cell-type) information present within the datasets. To better characterize single cell gene regulatory networks under different but related conditions we propose the joint estimation of multiple networks using multiview graph learning (mvGL). The proposed method is developed based on recent works in graph signal processing (GSP) for graph learning, where graph signals are assumed to be smooth over the unknown graph structure. Graphs corresponding to the different datasets are regularized to be similar to each other through a learned consensus graph. We further kernelize mvGL with the kernel selected to suit the structure of single cell data. An efficient algorithm based on prox-linear block coordinate descent is used to optimize mvGL. We study the performance of mvGL using synthetic data generated with a diverse set of parameters. We further show that mvGL successfully identifies well-established regulators in a mouse embryonic stem cell differentiation study and a cancer clinical study of medulloblastoma.

## 1. Introduction

Gene expression arises from an underlying network of regulatory interactions between transcription factors, co-factors and signaling molecules [49, 58]. Elucidating the topology of this underlying network is a major goal in modern computational biology, as it will allow researchers to discover pivotal factors that can help differentiate between the phenotypes of varying cell states [15, 41]. Advances in next generation sequencing technologies has made it possible to profile the transcriptomes of individual cells, hence capturing expression levels of thousands of genes at a cellular resolution. This has further enabled scientists to discover natural grouping among cells and define cell-types on the basis of the transcriptome [8, 19, 18]. The variation present between these cell-types holds the key to inferring how genes transcriptionally regulate each other and how their expressions and interactions change across cell subgroups. Accurate inference of cell-type specific GRNs will lead to a deeper understanding of subgroup specific trancriptional regulation, uncover mechanisms that occur when cellular processes are perturbed and enable identification of potential therapeutic targets for diseases [48, 29, 54]. Despite its advantages, single cell RNA sequencing (scRNA-seq) datasets is undermined by a series of technical limitations, such as drop-out events (expressed genes undetected by scRNA-seq) and a high level of noise, which makes single cell gene regulatory network inference a difficult problem [15, 6, 1]. Thus, given the high dimensionality and the complex nature of these datasets, there is a pressing need to develop specialized computational methods for the inference of cell-type specific GRN’s [7, 43].

In general, scRNA-seq datasets from a single experiment often contain cells belonging to different cellular subtypes [8, 19]. As the cells belonging to the different clusters have originated from the same tissue, their networks may be assumed to share some common structure. Independent network estimation for each subgroup does not allow for incorporation of shared structures between the subgroups. Thus, there is a need to develop joint network estimation strategies that can borrow information across subgroups while retaining between subgroup heterogeneity. Dozens of algorithms have been proposed for reconstruction of gene regulatory networks in scRNA-seq datasets [7, 43, 37]. However, most of these algorithms consider cells to be identically and independently distributed samples which violates the assumption of natural subgroups present within the data. Given our assumption of a clustered dataset, we should be able to apply these algorithms to estimate networks from each cell subgroup separately. However, this approach would not take the commonality across different cells within the subgroup which may lead to information loss. Multiple algorithms have been proposed for joint estimation of networks from high dimensional data. Most of these methods assume that the data has a Gaussian distribution. Seminal papers by [16, 10] paved the way for penalized estimation of multiple Gaussian graphical models (GGM), and demonstrated the use of lasso based penalty functions for better estimation of the sparsity structures across multiple groups. Very few methods have been proposed for joint estimation of multiple networks from scRNA-seq datasets. [35] developed PIPER, a penalized local Poisson graphical model (LPGM) [2] for joint estimation of multiple networks in scRNA-seq datasets. One of the main limitations of PIPER is that the Poisson distribution has one single parameter characterizing both the mean and the standard deviation. Single cell datasets would be much better characterized by a negative binomial distribution which has a separate dispersion parameter or a zero inflated negative binomial distribution which could account for the excess zeroes. To account for the non-gaussian nature of the scRNA-seq datasets, [55] proposed a modification of the joint Gaussian copula graphical model (JGCGM) based on the Gaussian copula transformation proposed in [26]. [11] proposed a three step hybrid joint estimation strategy that relies on (a) integrated application of a Bayesian zero inflated Poisson (ZIP) based model imputation strategy and single cell imputation technique McImpute [22, 33], (b) data Gaussianization [27] and eventually (c) joint estimation of a Gaussian graphical model [10]. Contrary to [35], the last two proposed approaches estimate graphical models for continuous data and rely on a data transformation step for making the data continuous.

Recent work in graph topology inference focuses on ideas developed in graph signal processing (GSP), which extends classical signal processing concepts to data defined on nodes of a graph, i.e. *graph signals* [51]. GSP based graph learning (GL) approaches infer the graph structure from the observed graph signals based on assumptions made about the relation between the signals and the unknown graph G [13]. These assumptions include smoothness, where graph signals are assumed to have low-frequency representation in graph frequency domain, and stationarity, where graph signals are assumed to be stationary with respect to G [31]. Due to explicit representation of graph signals in the graph frequency domain, GSP based GL has more flexibility in modeling signals compared to previous network inference methods, such as statistical models reviewed above for GRN inference. Therefore, various GL algorithms have been proposed and shown to perform well in different applications [12, 23, 50, 40]. However, most of this prior work focuses on learning a single graph structure and is not suitable for joint learning of multiple related graphs. Recently, a method for joint inference of multiple graphs from stationary graph signals is proposed in [36], where graphs are regularized to have similar sparsity patterns.

In this paper, we present a multiview Graph Learning (mvGL) algorithm for joint inference of graphs from multiple datasets. Based on our previous work [24], where smoothness of graph signals with respect to the underlying topology has been shown to perform well for GRN inference, mvGL learns multiple graphs by deriving an optimization problem using smoothness assumption. Graphs corresponding to the different datasets are ensured to be similar to each other through a learned consensus graph. Compared previous joint graph learning method [36], we focus on smoothness because of following reasons. First, smooth signals have low-frquency and sparse represenations in graph frequency domain, thus GL problem correspond to inferring efficient information processing transforms for graph signals. Second, many graph-based machine learning and signals processing tasks are developed based on the smoothness of the graph signals. Finally, many real-world applications are observed to have smooth signals [31]. mvGL is used for joint inference of GRNs across different cell types to unravel regulator genes whose interactions may vary across subgroups. During application to single cell data, mvGL is kernelized as in [24] to deal with the structure of scRNA-seq data. The proposed method has various unique advantages over existing approaches. First, it does not make any specific parametric assumptions about the data. Second, it employs specialized kernels to consider the nature of scRNA-seq data. For instance, it can employ proportionality measures as gene expressions in scRNA-seq reflect relative rather than absolute abundance. Similarly, zero-inflated Kendall’s tau can be used to handle drop-outs [52]. Finally, compared to [36], we focus onthe proposed method learns an additional consensus graph, which captures the common structure across all graphs.

## 2. Background

### 2.1. Graphs and Smooth Graph Signals

A weighted undirected graph can be denoted as G = (V, E) where V is the node set with |V| = n and E is the edge set. An edge between nodes i and j is represented by e_ij_ and it is associated with a positive weight w_ij_. Algebraically, a graph is represented by an adjacency matrix **W** *∈* **R**^n*×*n^ where W_ij_ = W_ji_ = w_ij_ if e_ij_ *∈* E and 0, otherwise. The combinatorial Laplacian matrix of G is defined as **L** = **D** − **W** where **D** is the diagonal matrix with node degrees, i.e. 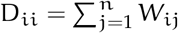. The Laplacian matrix is a positive semi-definite matrix, thus its eigendecomposition is **L** = **VΛV**^T^ where **Λ** is the diagonal matrix of eigenvalues. We assume the eigenvalues of **L** are ordered such that 0 = Λ_11_⩽ Λ_22_⩽ …, Λ_nn_. We represent vectors will all ones and zeros as **1**_n_ and **0**_n_, respectively, where the subscript indicates the size of the vector ^1^. The operator diag() : ℝ^n*×*n^ *→*ℝ^n^ returns the diagonal of the input matrix. The operator upper() : ℝ^n*×*n^ *→*ℝ^M^ returns upper triangular part of the input matrix where M = n(n − 1)/2. For an n *×* n symmetric matrix **A**, we define the matrix **S** *∈* ℝ^M*×*n^ such that **S**upper(**A**) = **A1** − diag(**A**).

A graph signal is a function x : V *→*ℝ and can be represented by a vector **x** *∈* ℝ^n^ where each x_i_ is the signal value on node i. Graph Fourier transform (GFT) of **x** can be defined using spectrum of **L** as its eigenvalues and eigenvectors provide a notion of frequency, i.e., small eigenvalues correspond to low frequency and larger ones to high frequencies [51]. GFT of **x** is then defined as 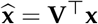 and inverse GFT is 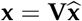. Thus, **x** is the linear combination of eigenvectors of **L** with the coefficients determined by 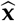. If most of the energy of 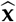 is concentrated in its first entries, **x** has a low-frequency representation in graph Fourier domain, i.e. it varies smoothly over the graph. The smoothness of **x** can be quantified by the total variation as:

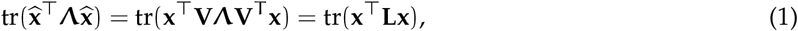

where smaller values indicate a smoother graph signal **x** with respect to G.

### 2.2 Single View Graph Learning

Given a set of graph signals 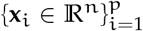 that are assumed to be smooth over an unknown graph G, the structure of G can be learned by the following optimization problem [12, 23]:

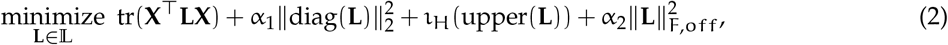

where **X** *∈* ℝ^n*×*p^ is the data matrix whose columns are **x**_i_’s, 𝕃 = {**L** : L_ij_ = L_ji_ ⩽0 ∀ i ≠ j, **L1** = **0**} is the set of valid Laplacian matrices. The first term in (2) measures the smoothness of the graph signals. The second term regularizes the degree distribution of the graph. The third term is the indicator function for hyperplane H = {**v** : **v**^T^**1** = −n} and is included to prevent the trivial solution **L** = **0**. The final term is the Frobenius norm of the off-diagonal entries of **L** and controls the density of the learned graph. Hereafter, the problem in (2) is referred to as single view graph learning (svGL).

## 3. Method

In this section, the proposed method to infer multiple GRNs from multiple cell types is described. The method is developed by deriving an optimization problem that extends svGL to handle multiple datasets. The optimization problem is then solved by prox-linear block coordinate descent (BCD) algorithm.

### 3.1 Multiview Graph Learning

Assume that we are given N datasets 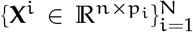 where the columns of **X**^i^ are p_i_ graph signals defined on an unknown graph G^i^ = (V, E^i^, W^i^). All G^i^’s have the same vertex set V with |V| = n. Their respective edge sets E^i^’s and associated edge weights W^i^ are assumed to be different, but similar to each other. Due to this assumption, inference of G^i^ can be improved by incorporating information from other graphs. For instance, in the analysis of scRNA-seq expressions, the datasets generated from a single tissue comprises of cells belonging to multiple cell-types. As cells having different cell-type assignments originate from a single tissue, their networks G^i^ may be considered to be related to each other. Learning G^i^’s independently from each cell-type does not allow for information borrowing from graphs of other cell types. Thus, the estimation of cell-type specific graphs can be improved by simultaneously learning multiple cell-type graphs. To this end, we propose an optimization problem, where we learn G^i^s simultaneously while regularizing them to be close to a consensus graph G. During the optimization, G is also learned by combining information from G^i^’s; thus G^i^’s are ensured to be similar to each other through G. Let **L**^i^’s and **L** be the Laplacian matrices of G^i^’s and G, respectively, then the corresponding optimization problem is:

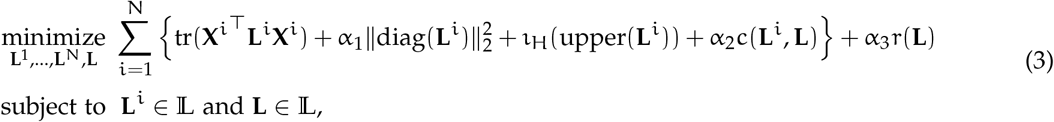

where the first term in the summation measures the smoothness of **X**^i^ over G^i^, the second and third terms have the same meaning as in svGL. The function c(**L**^i^, **L**) is included to ensure that each **L**^i^ is close to **L**. Finally, r(**L**) is a regularizer that controls the sparsity of **L**. While different choices for c(*·, ·*) and r(*·*) can be considered, in this work, 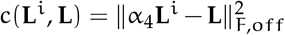 and r(**L**) = ‖**L**‖_1_ where α_4_ is a free variable introduced to control the sparsity of G^i^’s. (see Section 3.3 for further details about the selection of hyperparameters). Finally, (3) can be kernelized to exploit various (nonlinear) relations between graph signals. Kernelization is important for GRN inference as it is unclear which association measure between gene expressions is best performing for various scRNA-seq data analysis [52]. Therefore, following [24], we kernelize (3) by noting 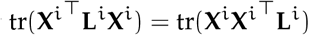 and changing 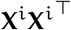 with any positive semi-definite kernel matrix **K**^i^. In this paper, we consider three kernels: correlation coefficient, r, proportionality measure ρ [45] and zero-inflated Kendall’s tau (τ_zi_) [42]. These kernels are considered because r is a commonly used for network inference, ρ is found as the best performing measure in [52] and τ_zi_ can handle of dropouts in scRNA-seq.

### 3.2. Optimization

The problem in (3) can be written in vectorized form, where one learns the upper triangular parts of **L**^i^’s and **L**. Let **k**^i^ = upper(**K**^i^), **d**^i^ = diag(**K**^i^), 𝓁^i^ = upper(**L**^i^) and 𝓁 = upper(**L**). The vectorized form of (3) is:

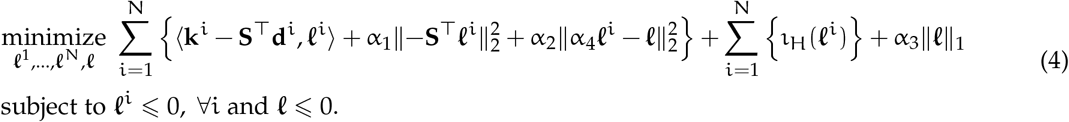

This problem can be solved using block coordinate descent (BCD) as its objective function is the sum of a smooth term and a separable non-smooth term. At each iteration of BCD, the objective function is minimized cyclically over each block, i.e. 𝓁^i^’s and 𝓁, while fixing the other blocks [56]. The subproblems considered at each iteration of BCD can be solved exactly or inexactly. When minimizing with respect to 𝓁^i^’s, we perform an inexact minimization using prox-linear update, as it results in an easy-to-solve subproblem and has fast convergence when extrapolation is employed. When minimizing with respect to 𝓁, we perform an exact minimization. More specifically, let F(𝓁^1^, …, 𝓁^N^, 𝓁) be the the first term in (4) and 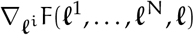 be its gradient with respect to 𝓁^i^. At iteration t, 𝓁^i^ is updated as follows:

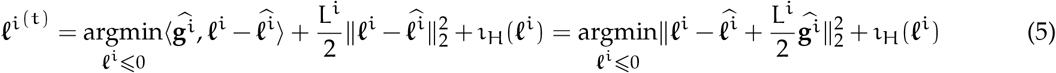

Where 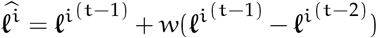 is the extrapolated point with w being the extrapolation weight, 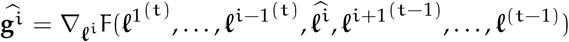 and L^i^ is the Lipschitz constant of 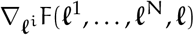. The solution of this problem is the projection of 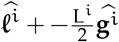 onto the negative simplex [14]. After all 𝓁^i^’s are updated, the update of 𝓁 is obtained by solving (4) with respect to 𝓁 :

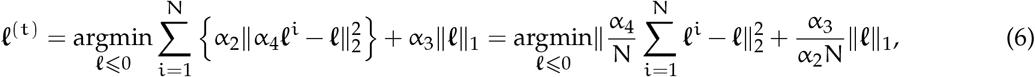

which is the proximal operator of 𝓁_1_ norm evaluated at 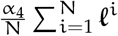 [39]. The convergence of this approach for solving (4) can be found in [56].

### 3.3. Hyperparameter Selection

There are four parameters that need to be selected for mvGL. α_1_ controls the degree distribution of the learned graphs such that larger values of α_1_ make all nodes have very similar degrees. Since degree distribution of real world networks is generally heterogeneous, we set α_1_ to the smallest possible value. However, if α_1_ is too small, the learned graph may become disconnected. Thus, we set α_1_ to the smallest possible value which ensures heterogeneous degree distribution and connectedness of the learned graphs. α_4_ controls the edge density of G^i^’s, i.e., larger values of α_4_ give denser graphs. We set it to a value that provides a desired edge density. Similarly, α_3_ controls the density of G such that as α_3_ gets larger G becomes sparser. Thus, we determine α_3_ in the same way as α_4_. Finally, α_2_ controls how close G^i^’s are to G, thus it can be set to a value which results in a desired correlation between G and G^i^’s. In the results section, the correlation between G and G^i^’s are determined so that there will be a weak correlation between G^i^’s, which means views are related to each other but view-specific features are also allowed to be learned.

## 4. Results

In this section, the performance of mvGL is evaluated on simulated data and two real scRNA-seq datasets. For simulated data, learned graphs are compared to ground truth networks to quantify the performance of mvGL. We employed area under precision recall curve (AUPRC) as the performance metric during this analysis. The performance of mvGL is benchmarked against three GRN algorithms and svGL on simulated datasets. We consider GENIE3, GRNBOOST2 and PIDC methods as these are found to be the best performing GRN inference algorithms for scRNA-seq data [43]. These methods and svGL can only learn a single graph from each dataset at a time^2^. Therefore, they are applied to each X^i^ separately and learned graphs are compared to ground truth G^i^’s. Additionally, they are applied to concatenation of X^i^ as a way to infer G, i.e. the consensus graph. Results for three different kernels are reported for mvGL and svGL: proportionality (ρ), correlation (r) and zero-inflated Kendall’s tau (τ_zi_). Hyperparameters of mvGL are set as described in Section 3.3. α_1_ and α_2_ for svGL control the degree distribution and density of the learned graph, respectively. Their values are set the same way as α_1_ and α_4_ are determined in mvGL.

For the analysis of real datasets, we first note that GRN can include both activating and inhibitory edges. In [24], it is observed that two genes that are connected by an activating edge have similar expressions. On the other hand, if they are connected with an inhibitory edge, their expressions are dissimilar. From GSP perspective, the former observation implies that the graph signal, i.e. gene expressions in a cell, has a low-frequency representation over the graph defined by activating edges, while the latter implies the graph signal has a high-frequency representation over the graph defined by inhibitory edges. mvGL as formulated in (3) only learns activating edges, because it assumes that graph signals have low-frequency representation over the unknown GRN. However, it can also be used to infer inhibitory edges. In particular, if the smoothness of the signals is maximized with respect to **L**^i^’s and **L**, the learned edges will be inhibitory. Thus, by changing **K**^i^ with −**K**^i^, we are able to learn inhibitory edges of the GRN. Based on this observation, results for both activating and inhibitory edges are reported in real data analysis.

### 4.1. Simulated Data

To validate the performance of mvGL, we simulate gene expression data from a multivariate zero-inflated negative binomial (ZINB) distribution. The ZINB distribution has been shown to accurately capture the characteristics of single cell datasets in several published studies [46, 17]. Given a known graph structure, we generate synthetic datasets using an algorithm developed by [57] and illustrated in [22, 24]. Two graph structures are considered for creating the baseline graph G: random graphs following an Erdős–Rényi model with an edge density of 0.1 and hub graphs following a Barabási–Albert model and having degree distribution that follows a power-law function. Since many real-world biological networks are known to possess scale free properties, hub networks present a more realistic representation of the single cell graph structures. Next, we generate K = 5 individual networks 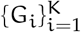 by adding ηM individual edges to the baseline graph G where M is the number of edges in G and η = 0.1. The ZINB simulator is then used to generate datasets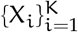, having n = 100 genes and p_i_ = 400 cells from the underlying graphs 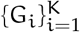. The three parameters of the ZINB distribution; λ, k and ω, which control its mean, dispersion and degree of zero-inflation, respectively were determined using a real scRNA-seq dataset [21]. This dataset has been previously used for estimating simulation parameters for a Bayesian zero inflated poison model in [11]. To investigate the effects of dropouts in graph reconstruction accuracy, we generate datasets by varying levels of zero-inflation between (0.26 − 0.36)%. Each simulated dataset include 10 random realizations and the average performance over realizations is reported.

Performance of methods on all datasets are shown in Figure 1. Since most of the GRN algorithms learn fully connected networks, we set desired density for mvGL and svGL to a large value (around 0.5). For views, average AUPRC over all views and realizations are reported. Irrespective of which kernel is employed, mvGL is performing better than svGL. This indicates that the proposed method shares valuable information across views, which improves the performance. Existing state of the art GRN algorithms are observed to perform poorly, which is inline with findings in previous joint GRN inference works [55]. When comparing the performance of different kernels, proportionality measure is found to be the best performing kernel. This finding is inline with [52], where different association metrics are compared based on their performance on various learning problems from scRNA-seq data. We also report results for how well consensus graphs match with the baseline graph G. As mentioned before, consensus graphs of benchmarking methods is found by concatenating X^i^’s. As concatenation leads to increased cell numbers, AUPRC values of benchmarking methods increase compared to values for views. Even in this case, mvGL emerges as the best performing algorithm. Finally, performance of most of the methods drops with increasing zero-inflation level (Although there are cases where this is not the case, we believe this is due to randomized nature of algorithms and datasets.) Performance drop in mvGL with increasing dropouts is less than the drop observed in other algorithms. This indicates sharing information across views may bring robustness to dropouts.

**Figure 1:**
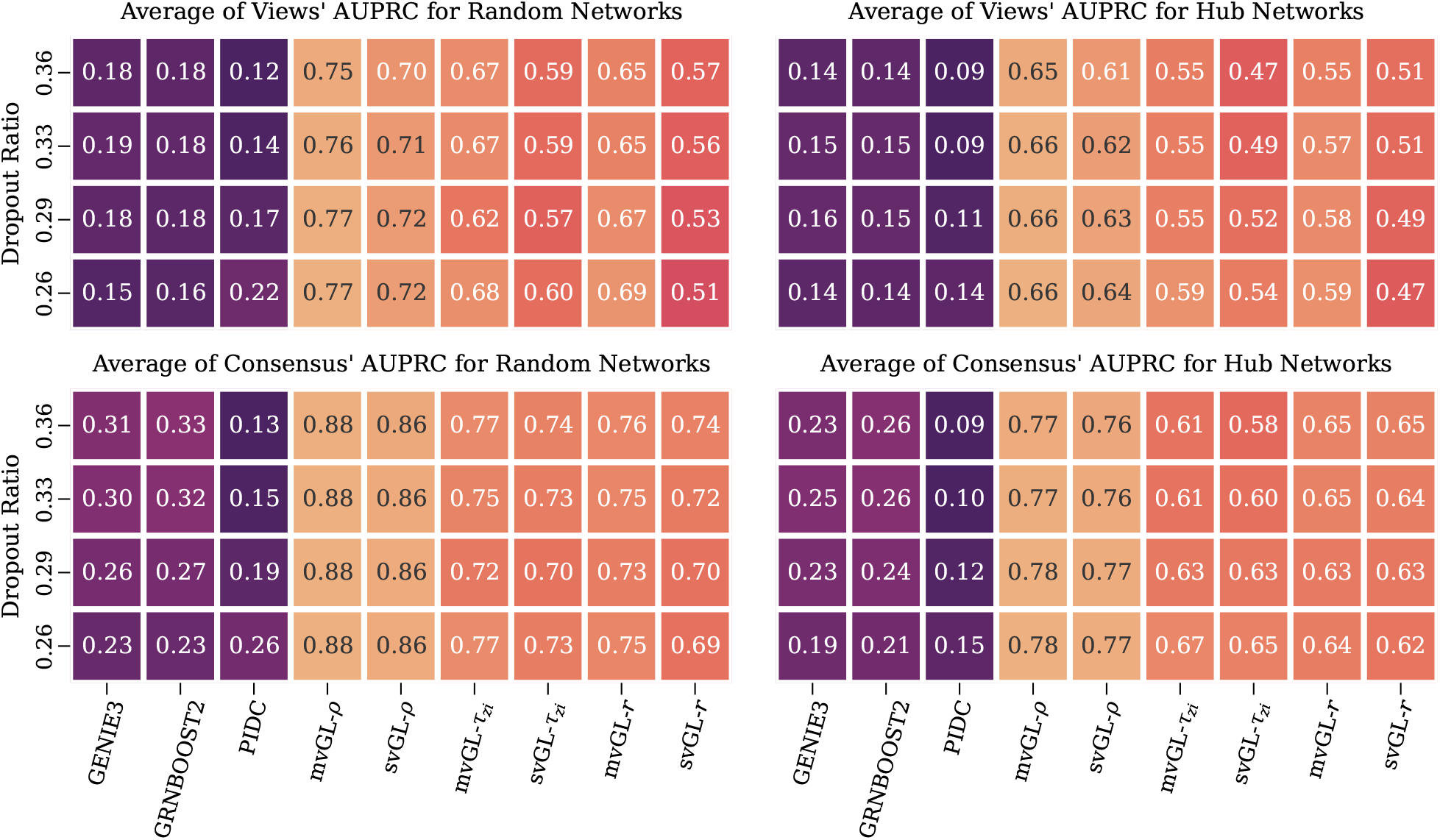
Performance of mvGL and svGL with three different kernels on simulated data. Top plots indicate AUPRC values for G^i^’s, where average over views and 10 realizations is reported. Bottom plots show AUPRC values for G, where average over realizations is reported.

### 4.2. Analysis of scRNA-seq data from mouse embryonic stem cell differentiation

Central to differentiation process and many other cellular processes is the expression of right combination of genes or modules of genes. Accurate characterization of the co-expression networks for progenitor and multiple cell types can help in understanding the cascade of cellular state transitions [32]. In this real data example, we study the differentiation process of mouse embryonic stem cells (mESC) using single cell RNA sequencing datasets [25]. This data was generated using high-throughput droplet-microfluidic approach and was primarily used to study differentiation in mESC before and after leukemia inhibitory factor (LIF) withdrawal. Since LIF maintains pluripotency of mESC, LIF withdrawal is considered to initiate the differentiation process. The dataset contains cells sampled from 4 states (or natural subgroups): before leukemia inhibitory factor (LIF) withdrawal, day 0 and after the withdrawal LIF for 2, 4, 7 days respectively. The subgroups contain 933, 303, 683 and 798 cells, respectively. This dataset has been previously analysed for studying joint graphical estimation in [35, 55] and similar to them we only consider the 72 stem cell markers for our example [44].

As per our workflow, we first estimated the subgroup specific and the consensus graphs, which are learned with activating and inhibitory edge densities set to 0.1. Based on the results of simulated data, we study graphs learned by proportionality measure. Next, we learned and ranked the hub genes for each of the learned graphs. Since gene networks are known to contain a handful of hub nodes that play very important roles via their many diverse connections, analysis of these hub nodes should highlight genes that are key to maintaining the cell differentiation process [3]. Figure 2 reports the hub genes learnt by mvGL. Our analysis confirms the importance of regulator genes Nanog, Sox2, Pou51, Zfp42, Utf1 in early stages of differentiation. Sox2, Nanog and Pou51 are known to play a fundamental role in the self-renewal and pluripotency of mouse embryonic stem cells [59] and therefore have been correctly identified as hub genes in the early stages of differentiation. Zfp42 and Utf1, both markers of pluripotency [30, 53] are also correctly identified as hub genes in Day 0 and Day 2 respectively. Reduction in expression of Nanog has been shown to be correlated with the induction of genes GATA4 which initiates differentiation of pluripotent cells [20] and therefore it has been correctly identified as hub genes in Day 2.

**Figure 2:**
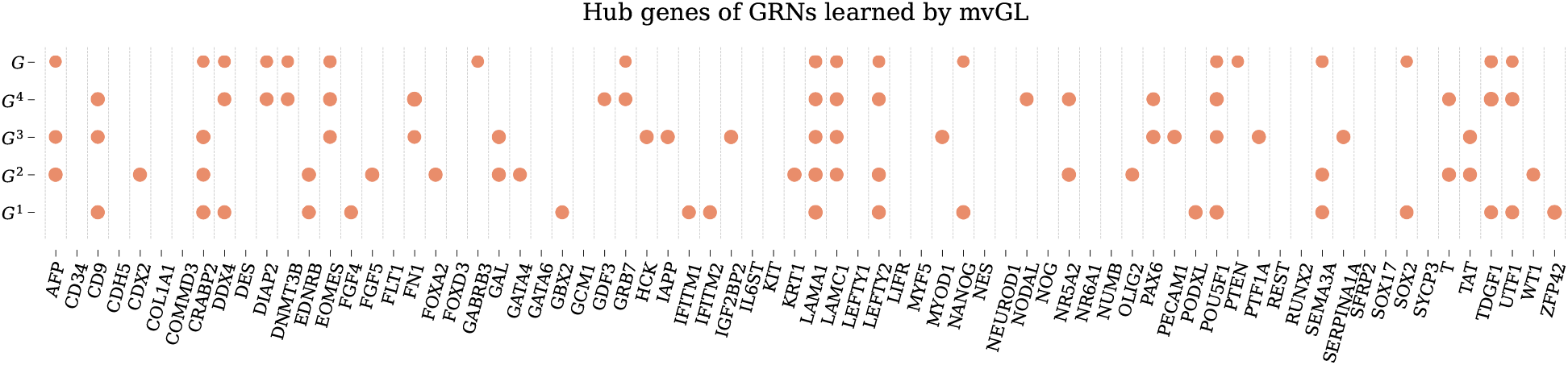
Hub genes in learned GRNs by mvGL for mESC dataset. To determine hub genes, degrees of the genes are sorted in descending order and the genes in the top-25th percentile are identified as hub genes. G^1^, G^2^, G^3^ and G^4^ correspond to Day 0, Day 2, Day 4 and Day 7, respectively.

### 4.3. Analysis of scRNA-seq data from medulloblastoma

Medulloblastoma (MB) is a highly malignant cerebellar tumor mostly affecting young children [38]. Several studies have been done to pinpoint the genetic drivers in each of its four distinct tumour subgroups: WNT-pathway-activated, SHH-pathway-activated, and the less-well-characterized Group 3 and Group 4 [38]. Among these subgroups, Group 3 and Group 4 tumors account for majority of the MB diagnoses, with Group 3 MB having a metastatic diagnosis of approximately 50%. Transcription factors(TFs) Myc2 and Otx2 have commonly been identified as key oncogenic TFs in Group 3 and 4 tumorogenesis. [13] used their proposed joint single cell network algorithm to study the roles of Myc and Otx2 utilizing the MB scRNA-seq data set (GSE119926) by [21]. Using the same selected samples from a subset of 17 individuals that were grouped into three subsets Group 3, Group 4 and an intermediate cell type, we estimate the joint gene regulatory network for the three groups for ∼ 750 genes among which most are enzyme-related genes from mammalian metabolic enzyme database [9].

Subgroup specific networks along with the consensus graph was estimated using proportionality kernel. The desired activating and inhibitory edge densities are set to 0.1. Figure 3 shows that Myc had a higher degree density for Group 3 in comparison to Group 4 and the intermediate subgroup. This confirms the major role MYC plays in initiation, maintenance, and progression in Group 3 tumors [47]. ALDH3A2 and ALDH1A2, the top ranking hub nodes were found to be downregulated in all the tumor subgroups confirming their role in cancer resistance [34, 5]. mvGL was also able to detect relationships between Myc and metabolic genes PAICS, PPAT, IMPDH1, ATIC and GART only in Group 3 tumors, however they did not make it to the top 25% of the MYC connections reported in Figure 3. These genes related to the human purine biosynthesis pathways have been previously reported to be induced and directly bound by Myc [28]. PRPS2, which emerged as one of strongest hubs in the OTX2 network analysis been known to promote increased nucleotide biosynthesis in Myc-transformed cells. This confirms that OtX2 is functionally cooperating with Myc to regulate gene expression in medulloblastoma [47]. mvGL also reported unique sets of hub genes for the OTX2 network confirming the unique role of OTX2 in MB tumorogenesis [11]. All of the results obtained indicate that mvGL was able to correctly identify the important functional interaction between OTX2 and MYC in regulating gene expression in medulloblastoma [4].

**Figure 3:**
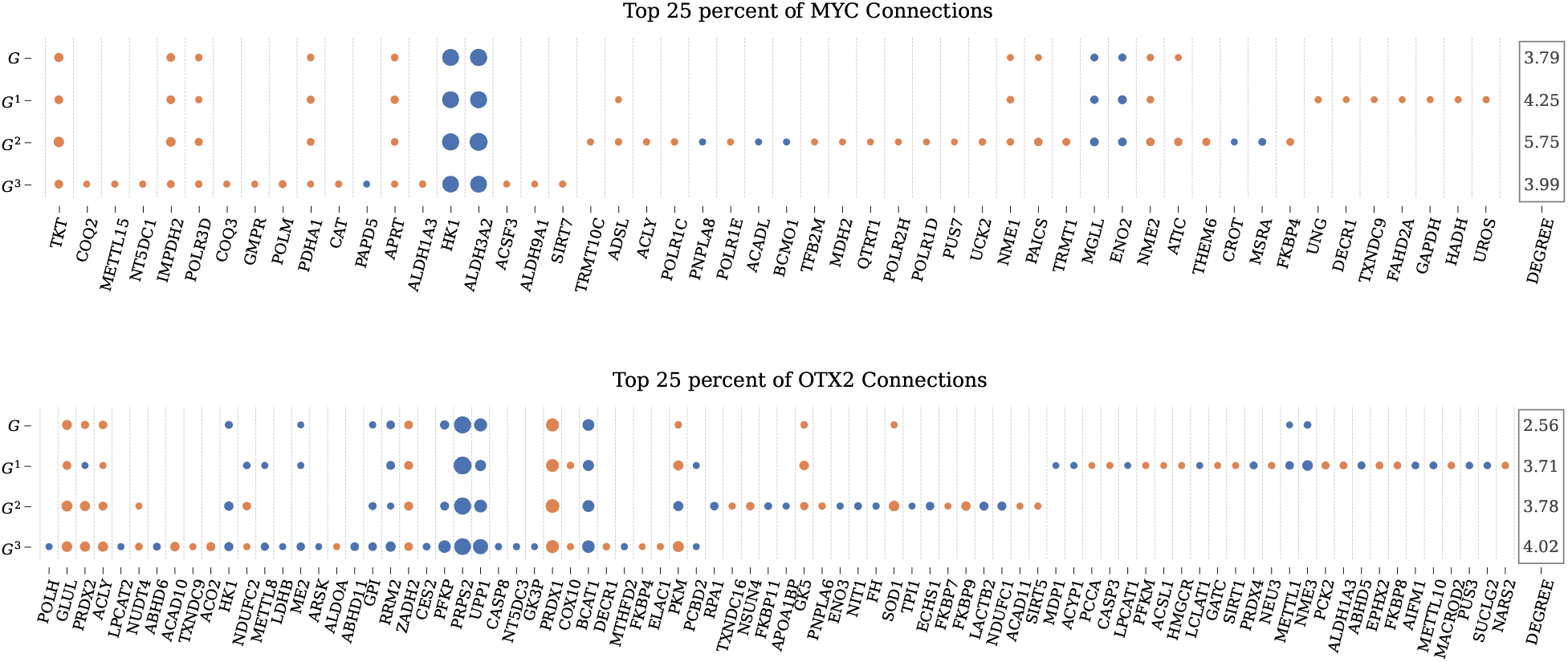
Neighbors of MYC and OTX2 genes. Marker size is proportional to connection weights. Orange and blue indicate that the connection is activating and inhibitory, respectively. Only connections that fall into top 25 percentile are reported. G^1^, G^2^ and G^3^ correspond to Intermediate, Group 3 and Group 4, respectively. At the right most part of the figure, total degrees of two genes in each graph are also shown.

## 5. Conclusion

In this paper we presented a multiview Graph Learning (mvGL) algorithm for joint inference of graphs from multiple scRNA-seq datasets. mvGL, a robust non-parametric approach, borrows ideas from kernel based graph signal processing for estimating the sparse high-dimensional structure of the underlying gene regulatory networks. Using simulation studies, we demonstrated the superior performance of mvGL over single view learning for random and hub graph topologies. Additionally the performance of mvGL remained robust to increase in dropout levels. We further analyzed two real single cell datasets and mvGL was shown to identify the key regulators in each of the differentiation and disease studies.

Future work can be aimed at inferring the multiple graph structures with a combination of kernel choices. That would ease the step of choosing a kernel suited to the particular dataset. Another limitations of mvGL is hyperparameter selection, which remains as an open problem in graph learning literature. In this work, we set them by manually searching hyperparameter space for values that provide graphs with desired densities and a desired correlation across views. Since, there is a monotonic relation between α_3_ (α_4_) and edge densities of the learned graph, this search can be performed automatically, which we leave as a future work. Incorporating combinational kernel strategies with improved hyperparameter learning could potentially further improve the accuracy of the estimated graphs.

## Footnotes

1 The subscript is ignored for the notational clarity in case the size is deducible from the context.

2 Joint GRN inference algorithms are fairly new and their toolboxes are not publicly available. Therefore, we could only benchmark against the methods that learn from single dataset.

